# Dietary vitamin B5 positively regulates intestinal neuropeptide secretion in *C. elegans*

**DOI:** 10.64898/2026.10.13.758927

**Authors:** Qi Jia, Andrew Calof, Derek Sieburth

## Abstract

Vitamin B5 is an essential vitamin that must be acquired via diet or by the gut microbiota, and vitamin B5 deficiency or altered vitamin B5 metabolism leads to a variety of physiological and neuronal disorders. We conducted a genetic analysis of the effects of exogenous vitamin B5 in *C. elegans,* and we found that vitamin B5 confers protection to toxicity caused by oxidative stress and acute exposure to vitamin B5 positively regulates neuropeptide secretion. We identified a sodium-dependent multivitamin transporter family member, *smvt-1* that is required for B5 uptake into cells and for protection conferred by vitamin B5. Vitamin B5 positively regulates the intestinal secretion of FLP-2, a neuropeptide-like protein whose release activates the antioxidant response. Vitamin B5 promotes FLP-2 secretion rapidly, specifically, and in a manner that is dependent upon uptake into intestinal cells by *smvt-1*. Vitamin B5-induced neuropeptide secretion is strictly dependent upon the SNARE protein *aex-4*/SNAP-25 and on the calcium dependent protein kinase C family member, *pkc-2/*PKCa in the intestine. These results reveal a previously uncharacterized function for dietary vitamin B5 in positively regulating intestinal neuropeptide secretion from dense core vesicles and in shaping the antioxidant response.

## Introduction

Organismal health is maintained through symbiotic interactions between the host organism and the gut microbiome. The gut microbiome produces metabolites that can communicate with the host’s nervous system, forming a functional gut-brain axis that can modulate neuronal function and behavior [1]. One such metabolite produced by gut bacteria is vitamin B5, or pantothenic acid, which is a precursor of coenzyme A (CoA). CoA is a cofactor that plays critical roles in diverse biochemical processes including enzymatic regulation, energy production, fatty acid metabolism, and protection against oxidative stress [2–6]. As an essential vitamin, vitamin B5 cannot be synthesized by eukaryotic cells and is instead taken up into cells by the sodium dependent multivitamin transporter, SMVT. Deficiencies in dietary vitamin B5 or uptake can result in defects in energy production, epithelial function, and lipid sequestration [7, 8], and reduced vitamin B5 levels in the nervous system are associated with Alzheimer’s and Huntington’s Disease [9, 10]. After its transport into cells, vitamin B5 is enzymatically converted into a series of intermediate metabolites that serve as precursors for de novo CoA biosynthesis. Rare pathological variants in the genes encoding these enzymes are associated with cardiomyopathy or neurodegenerative disease arising from impaired CoA production [11–15].

A critical mechanism by which the gut communicates with distal organs, including the brain, is through inter-tissue neuropeptide signaling [16–21]. Neuropeptides are a diverse class of signaling molecules whose regulated release from neurons and non-neuronal tissues orchestrate organismal physiology and behavior. A number of metabolites generated by the gut microbiome can positively regulate the secretion of neuropeptide-like proteins from the intestine including short chain fatty acids, which trigger the release of the peptide GLP-1, and hydrogen sulfide, which modulates ghrelin secretion, from enteroendocrine L-cells [22, 23]. In *C. elegans*, neuropeptide-like proteins released from the intestine regulate a variety of processes, including stress responses, rhythmic behavior and fat accumulation [16, 24–27]. We previously identified a gut-to-neuron endocrine circuit whereby the FMRFamide-like peptide FLP-2 is secreted from the intestine in response to endogenous hydrogen peroxide potentiates a systemic oxidative stress response [28, 29]. Intestinal FLP-2 secretion is regulated by endogenously produced hydrogen peroxide, but whether FLP-2 secretion or the oxidative stress response is also regulated by metabolites produced by bacteria is unknown.

Here we report a novel function of vitamin B5 uptake into intestinal cells in protection from toxicity caused by oxidants and in positively regulating FLP-2 secretion from the intestine. We identify the sodium dependent multivitamin transporter homolog, *smvt-1*, as a vitamin B5 transporter, and we show that the effects of vitamin B5 on both oxidative stress resistance and neuropeptide secretion are positively regulated by *smvt-1* activity in the intestine. Vitamin B5 positively regulates FLP-2 secretion in a manner that is dependent upon the SNARE protein *aex-4/*SNAP25 as well as the protein kinase C homolog *pkc-*2/PKCa/b in the intestine. These results reveal a role for dietary vitamin B5 in activating the anti-oxidative stress response by potentiating an inter-tissue gut-to-nervous system signaling axis.

## Materials and methods

### Strains and transgenic lines

*C. elegans* strains were maintained on standard nematode growth medium (NGM) plates seeded with *E. coli* OP50 bacteria as food source at 20°C in the dark. Strains were synchronized by picking mid L4 stage animals and analyzed either immediately before treatment (for coelomocyte assay and intestine imaging) or as young adults 24hrs later (for juglone toxicity assay). The wild type reference strain was Bristol N2. The *smvt-1(vj413) X* strain was generated using CRISPR/Cas9-mediated gene editing, as previously described [30]. Mutants used in this study were outcrossed at least four times and listed in Supplementary Table 1.

Transgenic animals were generated by micro-injecting plasmid mixes into the gonads of young adult hermaphrodites following standard techniques as previously described [31]. Micro-injection mixes were prepared using expression constructs mixed with co-injection markers: pJQ70 (P*ofm-1::rfp,* 20ng/μL), pMH163 (P*odr-1::mCherry*, 40ng/μL) or pDS806 (P*myo-3::mCherry*, 20ng/μL) to a final concentration of 100ng/μL. At least three transgenic lines were examined for each transgene, and one representative line was used for quantification. Transgenic lines used in this study are listed in the Supplementary Table 1.

### Molecular biology

All gene expression vectors were constructed with the backbone of pPD49.26 (for intestinal expression), or pPD95.75 (for pan neuronal expression) and subsequently sequenced. A 1.5kb fragment before the start codon of *rab-3* was used for pan-neuronal expression [32], a 2.0kb fragment before the start codon of *ges-1* was used for intestinal expression [33]. Promoter fragments were amplified from genomic DNA and inserted into the vector using *SphI* and *BamHI*. The *smvt-1* cDNA fragment (1.7kb) was amplified from a cDNA library and inserted into the vector using *NheI* and *KpnI*. For *smvt-1* localization, a *venus* fragment was ligated between the 13^th^ transmembrane domain and cytosolic tail of SMVT-1 using *AgeI* and *AvrII* and the *smvt-1::venus-DTLA* chimeric fragment was inserted into the vector using *NheI* and *KpnI*. Plasmids and oligos used in this study are listed in the Supplementary Table 1.

### CRISPR/Cas9-mediated editing

*smvt-1 (vj413)* deletion was created using a co-CRISPR protocol [30]. A single guide RNA (sgRNA) and a repair single-stranded oligodeoxynucleotides (ssODN) targeting *dpy-10* were co-injected with a sgRNA for genes of interest and a ssODN that induces homology-directed repair (HDR) to introduce Cas9-mediated mutagenesis. Cas9 guide RNAs were designed using the Alt-R^tm^ HDR Design Tool (IDT) and synthesized by IDTDNA targeting: 100bp before the start codon of *smvt-1* gene (5’-CCCAGTGACATCATTTCAAAACA-3’), exon 13 of *smvt-1* (5’-CCATTTCTTATCAGTATTACAGC-3’). ssODNs were ordered from Thermo Fisher Scientific. Fifteen young adult animals were injected to produce around thirty singled F1 animals carrying Dpy or Rol phenotype. F2 animals were genotyped for mutations based on PCR and enzyme digest. Homozygous mutants were outcrossed with wild type animals at least 2 times before being used for assays.

### Toxicity assay

Between 60-80 L4 stage animals were transferred onto a fresh NGM plate with OP50 and kept at 20°C in the dark overnight; the next day, a stock solution of 200mM juglone in dimethyl sulfoxide (DMSO) was freshly prepared and diluted into working solution of juglone with M9 buffer before treatment, adult animals were washed into a glass culture tube with juglone working solution and washed three times, then incubated at 20°C in the dark on a rotating mixer before being transferred onto a fresh NGM plate with OP50 to recover at 20°C in the dark overnight. Toxicity assays were performed in triplicate. Percentage of survival of assayed by counting the number of alive and dead animals.

### Microscopy and analysis

For coelomocyte imaging, L4 stage animals were transferred in fresh M9 buffer or working solution of juglone (diluted from a freshly prepared 200mM stock solution of juglone in DMSO) on a cover slide and washed six times. Animals were incubated for 10min before paralyzed with 30 mg/mL 2,3-butanedione monoxime (BDM) in M9 buffer. Images of coelomocytes next to the posterior end of the intestine were collected using the ×100 oil objective and average fluorescence intensity of Venus from the endocytic compartments in the posterior coelomocytes were measured in ImageJ.

For fluorescence distribution imaging, L4 stage animals were paralyzed with 30mg/mL BDM in M9 buffer and images of posterior intestinal region were collected using the GFP (excitation/emission: 500nm/520nm) and CFP (excitation/emission: 400nm/480nm) filter sets sequentially or as indicated.

Images were captured using the Nikon eclipse 90i microscope equipped with Nikon Plan Apo ×20, ×40, ×60, and ×100 oil objective (NA=1.40), and a Photometrics Coolsnap ES2 camera or a Hamamatsu Orca Flash LT+CMOS camera. Metamorph 7.0 software (Universal Imaging/Molecular Devices) was used to capture serial image stacks and to obtain the maximum intensity projection image for analysis.

### Liquid chromatography-tandem mass spectrometry (LC-MS/MS)

Worms were washed off NGM plates with M9 buffer and collected as described previously [34]. Analysis of relative quantities of vitamin B5 in *C. elegans* was conducted in biological triplicate (supplementary Table 2). To each tube containing a pellet of ∼1,000 animals, 200 µL of aqueous ascorbic acid (0.02 µg/µL) and ceramic beads were added. An Omni Bead Homogenizer was used to homogenize the C. elegans pellet. A BCA was conducted on a small aliquot from each group of harvested animals to determine protein concentration for normalized reconstitution before analysis. 100 µL of the sample was taken into a separate microcentrifuge tube, where 1 mL of methanol was added, and the tube was then mixed thoroughly. After centrifugation at 16,000 × g for 5 min, the supernatant was collected and dried in a vacuum concentrator. To each tube, phosphate buffer (500 μL, 500 mM K3PO4 (Sigma-Aldrich, cat # P5629) containing 4 mM Na2EDTA (Thermo Fisher Scientific, cat # 15576028), pH 8.5), dimethylaminopyridine (DMAP, 10 μL, 10 mM in acetonitrile) (Sigma-Aldrich, cat # 107700), and propionic anhydride (50 μL, 25% solution in acetonitrile) (Sigma-Aldrich, cat # 240311) were added and left to incubate for 15 min at room temperature. After drying the tubes in a vacuum concentrator, the samples are redissolved in a volume of 85/15 v/v water/acetonitrile, normalized to the BCA results, and vortexed for 5 min. After centrifugation at 16,000 × g for 5 min, the supernatant was collected and transferred to autosampler vials for subsequent analysis via LC-MS/MS.

A quantitative pantothenate assay was performed. A 10 uL aliquot of each sample was injected onto a GL Sciences InertSustain AQ-C18 (2.1×100 mm, 2.7μm, Agilent, Santa Clara CA) using a 1290 Infinity HPLC system (Agilent Technologies) and eluted at a flow rate of 300 µL/min with solvent A (100/0.1 water/formic acid) and solvent B (100/0.1 acetonitrile/formic acid) using an optimized 35 minute gradient (min/%B was: 0/15, 5/15, 23/50, 25/90, 27/15, 35/15). The column effluent was directed to an OptiFlow Turbo V Electrospray ionization source connected to a QTRAP 6500+ mass spectrometer (AB Sciex LLC), acquiring mass spectra in a targeted multiple reaction monitoring (MRM) mode in positive polarity. In this assay, detection of fragmented ions originating from each compound at specific LC retention times was utilized to ensure specificity and accurate quantification in the biological samples. Source parameters were as follows: Curtain Gas (CUR) 25 psi, Collision Gas (CAD) 9, IonSpray Voltage (IS) 4500V, Temperature (TEM) 550°C, and Ion Source Gases 1 and 2 (GS1, GS2) both 40 psi.

Chromatographic peak areas from the raw mass spectrometry data was integrated and extracted using Analyst (AB Sciex). The relative Analyte Peak Areas are compared between the two sample groups. Vitamin B5 concentrations in *C. elegans* was calculated as follows: [B5] in *C. elegans*=B5 Total (µmol)/(worm count/10µL×500µL). Cellular [B5]=[B5] in *C. elegans* (µmol)/1.4×10^-3^ µL.

### Statistical analysis

Statistical analysis was performed on GraphPad Prism 9. Unpaired t test with two tails was used for two groups and one-way ANOVA with multiple comparison corrections was used for three or more groups to determine the statistical significance. Statistical details and n are specified in the Figure legends. All comparisons are conducted based on wild-type controls unless indicated by lines between genotypes. Bar graphs with plots were generated using GraphPad Prism 9.

## Results

### Vitamin B5 supplementation is protective during oxidative stress

To determine whether vitamin B5 regulates the antioxidant response, we examined the survival of animals treated with the mitochondrial toxin juglone. Juglone inhibits the electron transport chain [35, 36], increasing mitochondrial ROS levels and juglone treatment of *C. elegans* leads to toxicity [37]. Treatment of *C. elegans* with 3uM of juglone for four hours results in significant lethality 24 hours later (Fig. 1a, [28, 29]). We found that one hour exposure to vitamin B5 prior to juglone treatment increased the percentage of animals that survive from 55% to over 70% (Fig. 1a). Thus, vitamin B5 exposure is protective to the toxic effects of juglone.

**Fig. 1.**
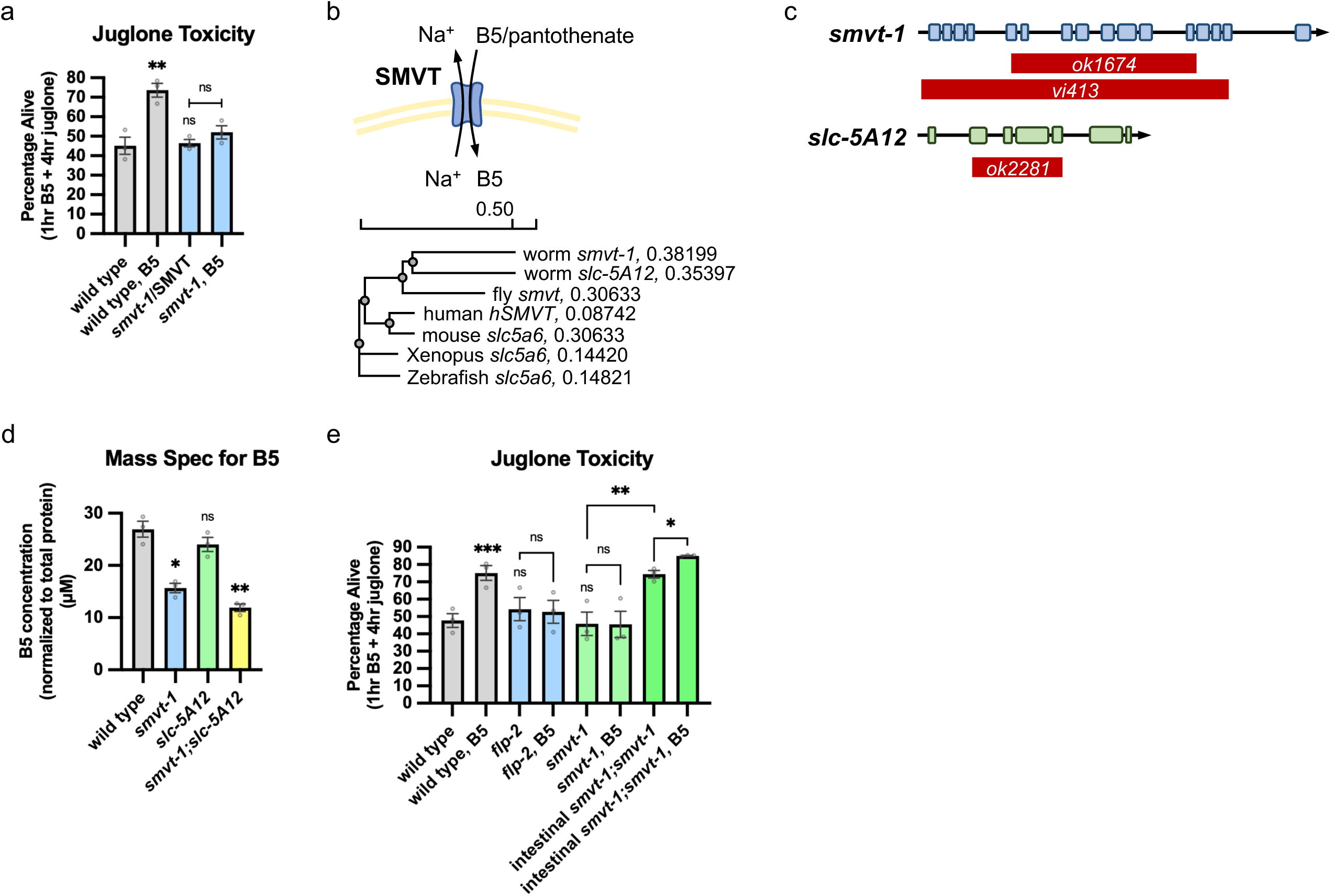
Vitamin B5 promotes organismal survival against juglone induced toxicity through its transporter SMVT-1 in the intestine. a) Average percentage of surviving young adult animals of the indicated genotypes after 16h recovery following 4h juglone treatment. Unlined ** and ns denote statistical significance compared to “wild type”. n = 268, 216, 167, 115, 150, 168, 255, 196 independent biological samples over three independent experiments. b) (Top) Schematic showing that sodium/multivitamin transporter (SMVT) actively transports pantothenic acid (vitamin B5) across cellular membranes in exchange for sodium. (Bottom) Phylogenetic tree showing relative distance of amino acid homology of SMVT family members across the indicated species. c) (Top) Schematic showing the positions of the deletions in the *ok1674* allele and *vj413* allele for *smvt-1* knockout. (Bottom) Schematic showing the position of the deletion in the *ok2281* allele for *slc-5A12* knockout. d) Average total vitamin B5 concentration calculated on a per cell basis of well-fed animals of the indicated genotypes normalized to total protein concentration. Vitamin B5 concentrations were based on a standard curve generated using purified pantothenate, and averages were calculated from biological triplicates. e) Average percentage of surviving young adult animals of the indicated genotypes after 16h recovery following 4h juglone treatment. Unlined *** and ns denote statistical significance compared to “wild type”. n = 210, 139, 277, 247, 215, 182, 180, 140 independent biological samples over three independent experiments. (a, d and e) Data are mean values ± s.e.m normalized to wild type controls. ns. not significant, * *P* < 0.05, ** *P* < 0.01, *** *P* < 0.001 by Brown-Forsythe and Welch ANOVA with Dunnett’s T3 multiple comparisons test.

### *smvt-1* encodes a vitamin B5 transporter

To determine whether vitamin B5 is taken into cells, we examined mutants lacking the putative vitamin B5 transporters. SLC5A6 encodes the mammalian sodium-dependent multivitamin transporter (SMVT), which mediates vitamin B5 uptake into cells in exchange for sodium ions (Fig. 1b). Human SMVT shares the highest amino acid homology with two *C. elegans* genes, *smvt-1* and *slc-5A12,* which are predicted to contain 12 transmembrane domains found in SMVT family members and are related to SMVT orthologs in a variety of organisms (Fig 1b).

To determine whether *smvt-1* and/or *slc-5A12* are vitamin B5 transporters, we first measured free vitamin B5 levels from lysates of well-fed *C. elegans* using quantitative liquid chromatography mass spectroscopy (LC-MS). Normalized to total protein concentration, wild type animals exhibited average total vitamin B5 levels of 3.772×10^-8^µmol per worm, which on a per cell basis is estimated to be 26.9µM, which is within the range of 1µM to 100µM reported for different mammalian cell types [38]. *smvt-1* putative null mutants harbor a deletion that removes nearly their entire coding region (Fig. 1c), and *smvt-1* mutants showed a 35% reduction in vitamin B5 levels compared to wild type controls (Fig. 1d). In contrast, *slc-5A12* putative null mutants, which harbor a deletion that removes nearly their entire coding region (Fig. 1c), had vitamin B5 levels that were similar to those of wild type controls (Fig. 1d). *smvt-1; slc-5A12* double mutants had a 53% reduction in vitamin B5 levels compared to wild type controls (Fig. 1d). Together these results indicate that both *smvt-1* and *slc-5A12* are likely to encode functional vitamin B5 transporters, and that *smvt-1* and *slc-5A12* function partially redundantly, with *smvt-1* being the primary vitamin B5 transporter and *slc-5A12* becoming critical in the absence of *smvt-1*. *smvt-1* and *slc-5A12* have distinct tissue expression patterns, with *smvt-1* expressed primarily in the nervous system and intestine, and *slc-5A12* expressed in muscle, skin and intestine [39]. Since vitamin B5 is still detected at low levels in the double mutants, vitamin B5 may also be transported into cells via SLC5A6-independent mechanisms, including by passive diffusion [40].

### *smvt-1* is required for the protective effects of vitamin B5

We next determined whether vitamin B5 uptake is required for its protective effects by examining survival of SMVT mutants. *smvt-*1 single mutants exhibited survival rates following juglone treatment that were not significantly different than wild type controls (Fig. 1e). However, the beneficial effect of vitamin B5 exposure on survival was abolished in *smvt-1* mutants (Figs. 1a and 1e). Expression of *smvt-1* cDNA selectively in the intestine (under the *ges-1* promoter) restored the protective effect of exogenous vitamin B5 to *smvt-1* mutants, and increased protection to oxidative stress-induced toxicity even in the absence of exogenously added vitamin B5 (Fig. 1e). These results indicate that exogenous vitamin B5 requires transport into intestinal cells by SMVT-1 for its protective effects against oxidative stress, and they suggest that in the absence of exogenous vitamin B5, SMVT-1 normally limits vitamin B5 transport into the intestine. Consistent with this, vertebrate SMVT has been shown to be limiting for vitamin B5 uptake in models of tumor growth [41].

### Vitamin B5 requires signaling by the neuropeptide-like protein *flp-2* for its protective effects

We previously showed that the antioxidant response is positively regulated by the neuropeptide like protein *flp-2* [29]. To determine whether the protective effect of exogenous vitamin B5 requires *flp-2* signaling, we examined *flp-2* null mutants. Mutations in *flp-2* abolished vitamin B5-induced protection against juglone-mediated toxicity (Fig. 1e). Thus, the *flp-2* signaling pathway functions downstream or in parallel to exogenous vitamin B5 to protect from juglone-mediated toxicity.

### Vitamin B5 rapidly positively regulates FLP-2 secretion from the intestine

We previously found that the secretion of FLP-2 from the intestine potentiates the antioxidant response and FLP-2 secretion is positively regulated by acute juglone exposure (Figs. 2a and b and [29]). To determine whether vitamin B5 can regulate FLP-2 secretion, we examined transgenic animals expressing FLP-2 fusion proteins tagged with Venus (FLP-2::Venus) selectively in the intestine. FLP-2::Venus fusion proteins secreted from the intestine are taken up by coelomocytes (scavenger cells) by bulk endocytosis, and the fluorescence intensity in coelomocytes is a measure of the efficacy of FLP-2 secretion (Fig. 2a and [29]).

**Fig. 2.**
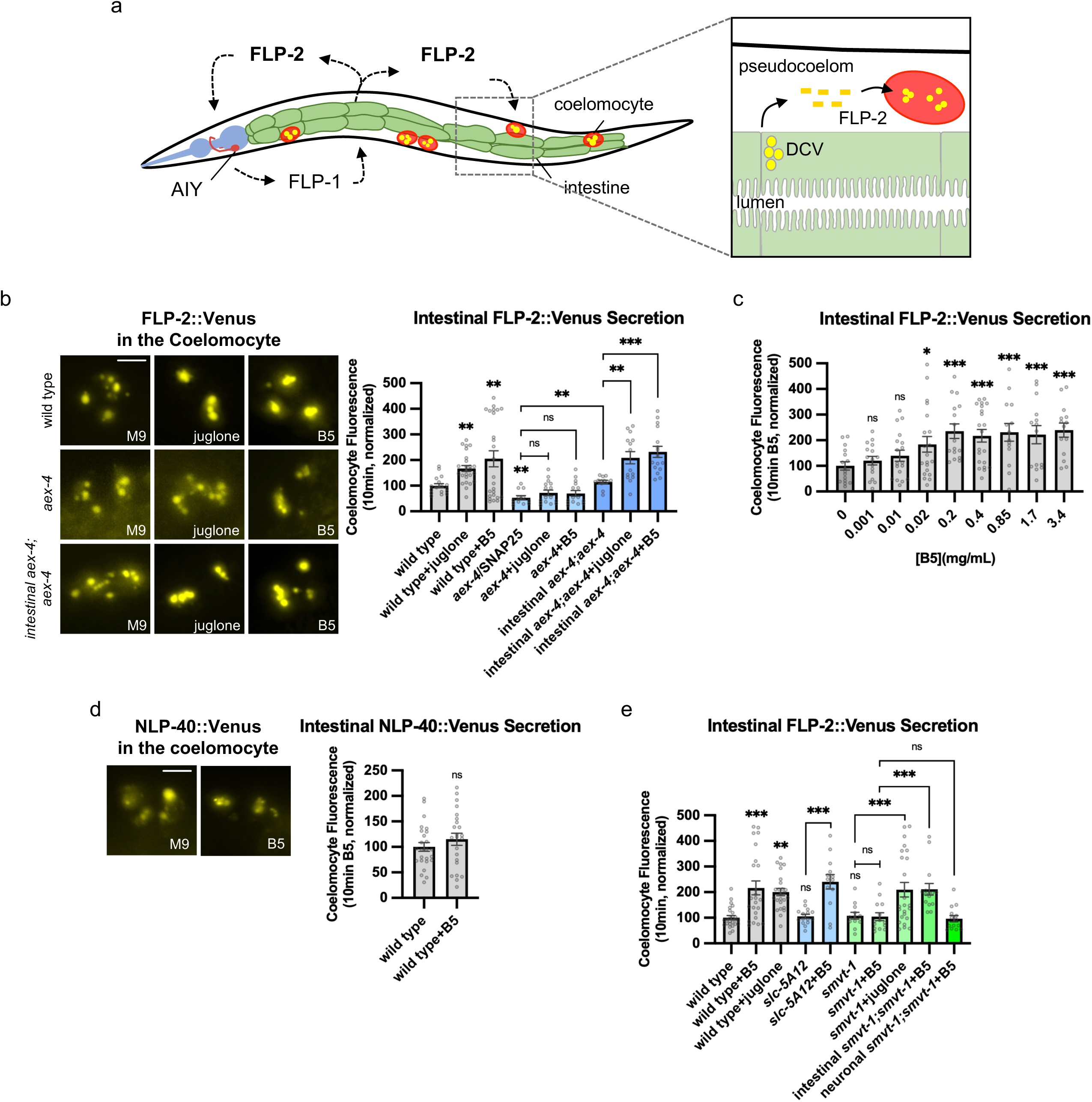
Vitamin B5 positively regulates the release of FLP-2 peptides from the intestine through SMVT-1. a) Schematic showing that FLP-2 released from the intestine activates AVL during the oxidative stress response. Once secreted, FLP-2 is also taken up by scavenger cells (coelomocytes). Fluorescence intensity of coelomocytes from transgenic animals expressing FLP-2::Venus in the intestine is used as an assay to measure FLP-2 secretion efficacy. b) Representative images and quantification of average coelomocyte fluorescence of the indicated mutants expressing FLP-2::Venus fusion proteins in the intestine following M9 buffer, 1mM juglone or 0.2mg/mL vitamin B5 for 10min. Unlined ** denotes statistical significance compared to “wild type”. n = 17, 25, 25, 12, 16, 15, 12, 14, 16 independent animals. Scale bar: 5μM. c) Quantification of average coelomocyte fluorescence of wild type animals expressing FLP-2::Venus fusion proteins in the intestine following M9 buffer or vitamin B5 in indicated concentration for 10min. Statistical significance was compared to “0mg/mL B5” (M9 buffer). n = 16, 16, 18, 20, 16, 21, 14, 14, 14 independent animals. d) Representative images and quantification of average coelomocyte fluorescence of wild type animals expressing NLP-40::Venus fusion proteins in the intestine following M9 buffer or vitamin B5 for 10min. n = 25, 23 independent animals. Scale bar: 5μM. e) Quantification of average coelomocyte fluorescence of the indicated mutants expressing FLP-2::Venus fusion proteins in the intestine following M9 buffer, 1mM juglone or 0.2mg/mL vitamin B5 for 10min. Unlined ***, ** and ns denote statistical significance compared to “wild type”. n = 25, 24, 25, 12, 14, 24, 15, 13, 15, 14 independent animals. (b-e) Data are mean values ± s.e.m normalized to wild type controls. ns. not significant, * *P* < 0.05, ** *P* < 0.01, *** *P* < 0.001 by Brown-Forsythe and Welch ANOVA with Dunnett’s T3 multiple comparisons test.

FLP-2::Venus-expressing animals exhibit low levels of coelomocyte fluorescence under normal conditions, indicating that there is some baseline FLP-2::Venus secretion (Fig. 2b). We found that a ten-minute exposure to vitamin B5 significantly increased coelomocyte fluorescence by about two-fold (Fig. 2b). This increase was similar to the increase elicited by ten-minute juglone exposure (Fig. 2b and [29]). The increase in coelomocyte fluorescence was dose-dependent, reaching a two-fold increase in fluorescence at 0.2mg/ml vitamin B5 concentrations that plateaued at higher concentrations tested (Fig. 2c). Vitamin B5 exposure had no detectable effect on the intestinal secretion of NLP-40 (Fig. 2d), which is neuropeptide-like protein that is secreted from the intestine in a juglone-independent manner [24, 29]. Neuropeptide secretion from the intestine requires *aex-4*/SNAP25, which encodes a t-SNARE protein that mediates the fusion of secretory vesicles with the plasma membrane, and the subsequent exocytosis of secretory vesicle cargo [29, 42]. Mutants lacking *aex-4*/SNAP25 have reduced coelomocyte fluorescence of FLP-2::Venus compared to wild type controls, and they exhibit no increase in coelomocyte fluorescence following juglone treatment (Fig. 2b and [29]), indicating that both baseline and juglone-induced FLP-2 secretion is dependent upon SNAP25-mediated DCV fusion. Similarly, vitamin B5-induced FLP-2 secretion was also dependent upon *aex-4*/SNAP25 since *aex-4* mutants treated with vitamin B5 failed to exhibit increased coelomocyte fluorescence (Fig. 2b). Expression of *aex-4* cDNA specifically in the intestine fully restored both baseline and vitamin B5-induced FLP-2 secretion to *aex-4* mutants (Fig. 2a). Together these results suggest that acute exposure to exogenous vitamin B5 selectively increases SNARE-mediated FLP-2 secretion form dense core vesicles in the intestine.

### Intestinal *smvt-1* is required for vitamin B5-induced FLP-2 release

To determine whether the cellular uptake of exogenous vitamin B5 is necessary to increase FLP-2 secretion, we examined *smvt-1* and *slc-5A12* mutants. We found that *smvt-1* mutants expressing intestinal FLP-2::Venus exhibited coelomocyte fluorescence that was similar to wild type controls (Fig. 2e), indicating that loss of *smvt-1* does not impact baseline FLP-2 secretion. Similarly, *smvt-1* mutants exhibited normal juglone-induced FLP-2 secretion (Fig. 2e). However, *smvt-1* mutants showed no significant changes in FLP-2 secretion following vitamin B5 exposure compared to untreated *smvt-1* mutants (Fig. 2e). Expression of *smvt-1* cDNA in the intestine restored full vitamin B5 responsiveness to *smvt-1* mutants, whereas pan-neuronal expression of *smvt-1* cDNA (under the *rab-3* promoter) failed to rescue (Fig. 2e). *slc-5A12* mutants exhibited baseline and vitamin B5-induced increases in FLP-2 secretion that were not significantly different than wild type controls (Fig. 2e). Thus *smvt-1* is required cell autonomously in the intestine for vitamin B5-induced FLP-2 secretion. Together these results indicate that the transport of vitamin B5 into intestinal cells by SMVT-1 is essential for positively regulating FLP-2 secretion.

### SMVT-1 localizes to the plasma membrane of intestinal cells

Mammalian SMVT is localized to the apical surfaces of intestinal cells where it functions as a transporter for vitamin B5 [43, 44]. To investigate the localization of SMVT-1, we examined animals expressing functional SMVT-1 fusion proteins with GFP in intestinal cells. We inserted GFP internally four amino acids from the C-terminal cytosolic tail of SMVT-1 (SMVT-1::GFP-DTLA, Fig. 3a). SMVT-1::GFP-DTLA fusion proteins fully rescued the FLP-2::Venus secretion defects of *smvt-1* mutants (Fig. 3b). Intestinal cells are polarized with the apical membranes facing the intestinal lumen, and the basal and lateral membranes facing the internal fluid cavity (the pseudocoelom). SMVT-1 tagged with mTurquoise2 at the same position (SMVT-1::mTur2-DTLA) localized to all three intestinal membranes, with lower levels detected on the apical and basal membranes, and higher levels on the lateral membranes (Fig. 3c).

**Fig. 3.**
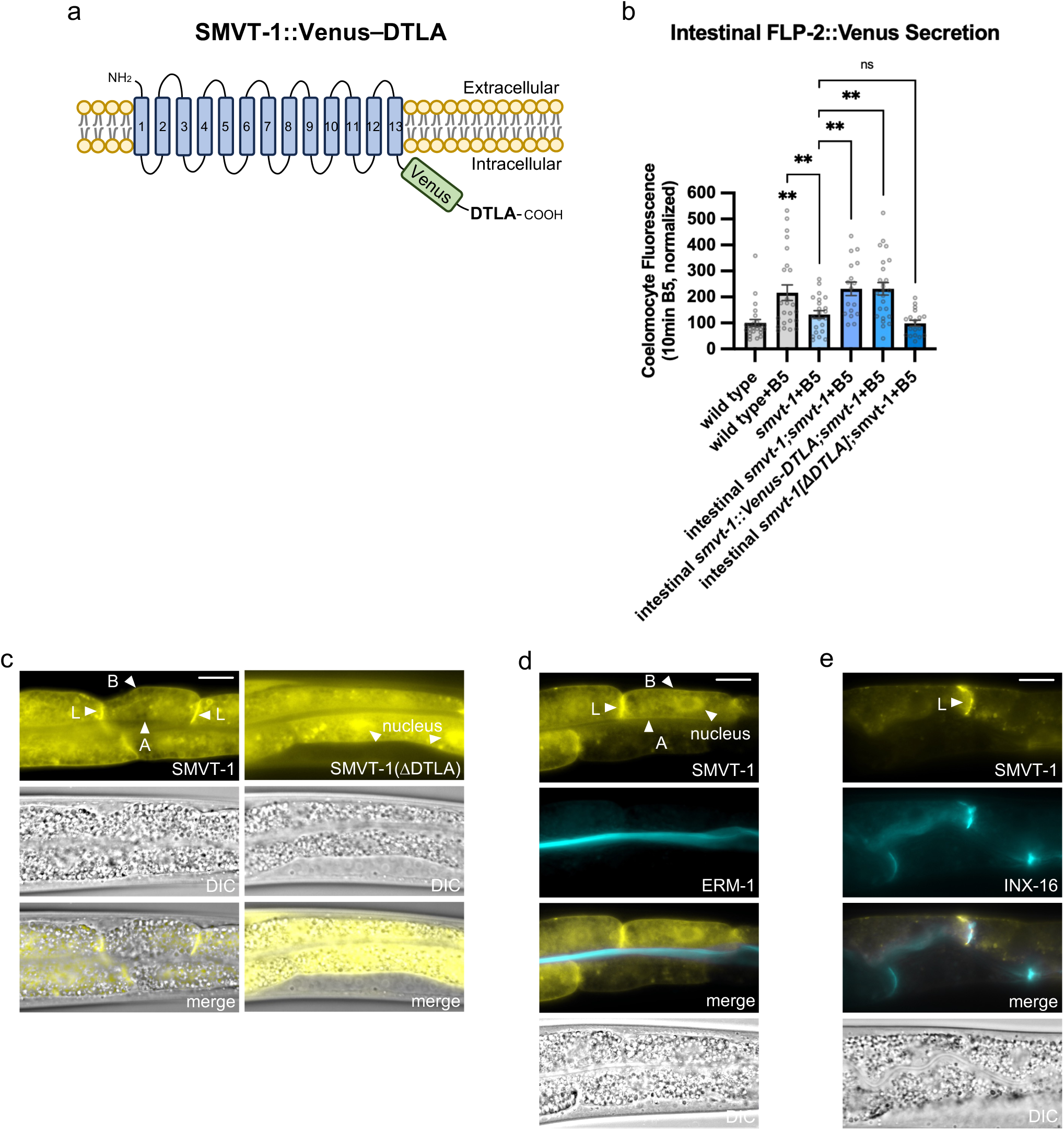
SMVT-1 localizes to the plasma membrane of intestinal cells. a) Schematic showing the design of SMVT-1::Venus-DTLA fusion protein and its localization on the plasma membrane. b) Quantification of average coelomocyte fluorescence of the indicated mutants expressing FLP-2::Venus fusion proteins in the intestine following M9 buffer or 0.2mg/mL vitamin B5 for 10min. Unlined ** denotes statistical significance compared to “wild type”. n = 25, 24, 22, 17, 25, 18 independent animals. Data are mean values ± s.e.m normalized to wild type controls. ns. not significant, ** *P* < 0.01 by Brown-Forsythe and Welch ANOVA with Dunnett’s T3 multiple comparisons test. c) Representative fluorescence images, merged images, and corresponding differential contrast images showing distribution in the posterior intestinal region of transgenic animals expressing SMVT-1::Venus-DTLA fusion proteins (Left) and SMVT-1::Venus (Right) in the intestine. (A), (B) and (L) mark the apical, basal and lateral membrane of the intestine cell. Scale bar: 20μM. d) Representative fluorescence images, merged images, and corresponding differential contrast images showing distribution in the posterior intestinal region of transgenic animals expressing SMVT-1::Venus-DTLA and ERM-1::mTur fusion proteins. (A), (B) and (L) mark the apical, basal and lateral membrane of the intestine cell. Scale bar: 10μM. e) Representative fluorescence images, merged images, and corresponding differential contrast images showing distribution in the posterior intestinal region of transgenic animals expressing SMVT-1::Venus-DTLA and INX-16::mTur fusion proteins. (A), (B) and (L) mark the apical, basal and lateral membrane of the intestine cell. Scale bar: 10μM.

The apical localization of SMVT-1 was confirmed with the apical intestinal marker ERM-1 [45] (Fig. 3d). The lateral intestinal membranes form gap junctions composed of INX-16/innexin with neighboring intestinal cells [46, 47], and the laterally-localized SMVT-1 colocalized with INX-16 (Fig. 3e). The apical localization of SMVT-1 is consistent with a role for SMVT-1 in transporting vitamin B5 from the intestinal lumen into the intestinal cells, while the basal and lateral localization implies that SMVT-1 may transport vitamin B5 from the pseudocoelom into intestinal cells or may represent non-functional pools of SMVT-1 that that are distributed to the apical membrane for vitamin B5 uptake.

The localization and function of mammalian SMVT is dependent upon a class 1 PDZ-binding motif (-ETSL) located at its C-terminus [43, 48]. Worm SMVT-1 contains a sequence at its C-terminus (-DTLA) that matches the class I PDZ-binding motif consensus -XS/TXΦ [48], where Φ is a hydrophobic amino acid. Deleting this motif of SMVT-1 resulted in a truncated protein (SMVT-1::[DDTLA]) that failed to rescue the FLP-2::Venus secretion defects of *smvt-1* mutants (Fig. 3b). Occluding the PDZ-binding domain by fusing GFP to the C-terminus of SMVT-1 (SMVT-1::GFP) resulted in a fusion protein that did not localize to the intestinal plasma membrane but instead adopted a cytosolic localization pattern (Fig. 3c). These results indicate that the PDZ binding motif of SMVT-1 is critical for both the plasma membrane localization and function of SMVT in the intestine, and they suggest that the molecular mechanisms underlying SMVT-1 plasma membrane targeting and/or retention may be mediated by evolutionarily conserved interactions between its C-terminus and PDZ domain-containing proteins.

### *pkc-2*/protein kinase C is required for the effects of vitamin B5 on FLP-2 secretion

Protein kinase Cs (PKCs) are serine threonine kinases that modulate regulated exocytosis in a wide variety of cell types [49]. We previously found that the calcium and diacylglycerol-stimulated PKC family member, *pkc-2/*PKCα/β, functions in the intestine for juglone-induced FLP-2 secretion [29]. To determine whether *pkc-2* is required for the effects of vitamin B5 on FLP-2 secretion, we examined *pkc-2* null mutants. *pkc-2* mutants exhibited basal secretion of FLP-2 that was similar to wild type controls. However, neither juglone nor vitamin B5 exposure could increase FLP-2::Venus secretion of *pkc-2* mutants above baseline levels (Fig. 4a). Expression of *pkc-2b* cDNA selectively in the intestine fully restored both juglone- and vitamin B5-responsivness to *pkc-2* mutants (Fig. 4a and [29]). These results show that vitamin B5-induced FLP-2 secretion is dependent on intestinal *pkc-2*, indicating that *pkc-2* may function downstream or in parallel to vitamin B5 to promote FLP-2 exocytosis.

**Fig. 4.**
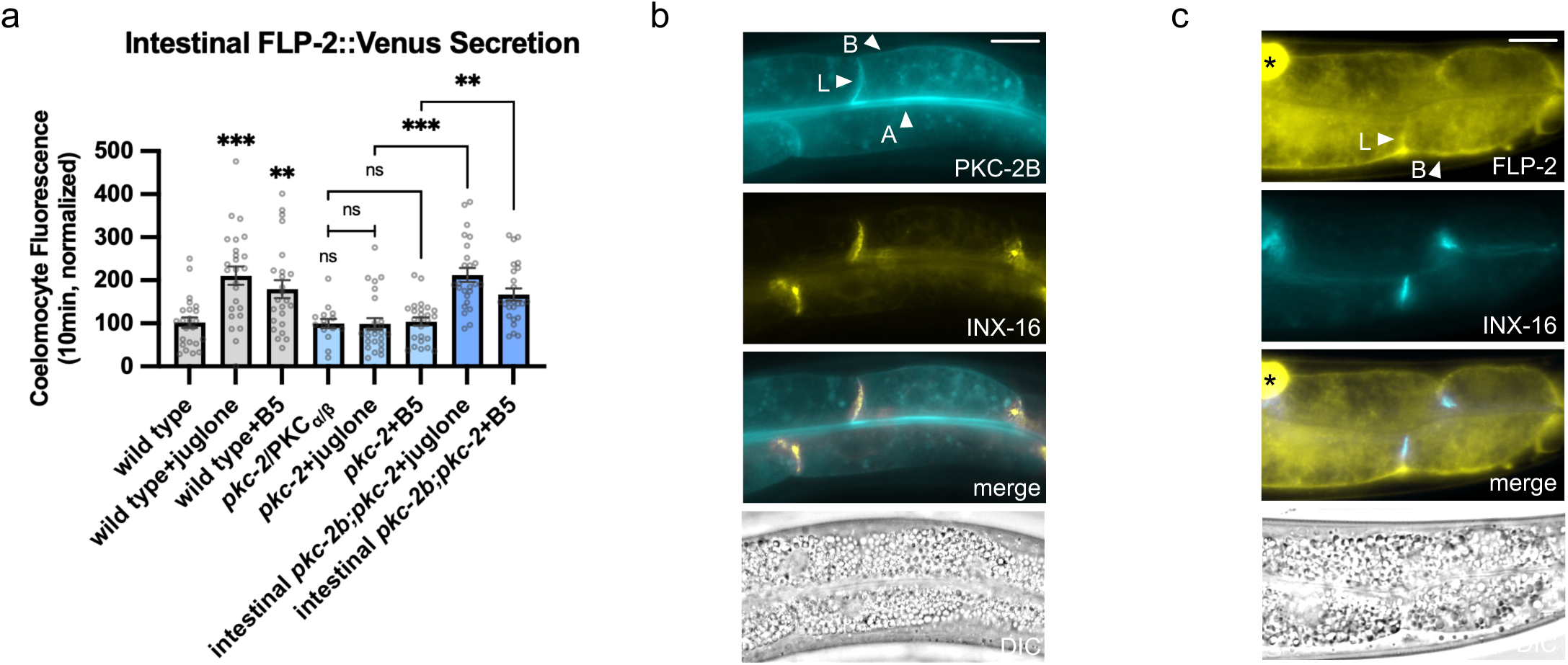
Vitamin B5 promotes FLP-2 release through protein kinase PKC-2. a) Quantification of average coelomocyte fluorescence of the indicated mutants expressing FLP-2::Venus fusion proteins in the intestine following M9 buffer, 1mM juglone or 0.2mg/mL vitamin B5 for 10min. Unlined ***, ** and ns denote statistical significance compared to “wild type”. n = 24, 25, 24, 15, 25, 25, 25, 25 independent animals. b) Representative images of fluorescence distribution in the posterior intestinal region of transgenic animals co-expressing PKC-2B::mTurquoise2 and INX-16::Venus fusion proteins in the intestine. Scale bar: 10μM. c) Representative fluorescence images, merged images, and corresponding differential contrast images showing distribution in the posterior intestinal region of FLP-2::Venus and INX-16::mTur fusion proteins. (B) and (L) mark the basal and lateral membrane of the intestine cell. Scale bar: 10μM.

### PKC-2 and FLP-2 localize to basal and lateral intestinal membranes

The activation of PKC members involve their translocation from cytosolic pools to cellular membranes via interactions with calcium and diacyl glycerol (DAG) [50]. To determine the localization pattern of PKC-2 in the intestine, we generated PKC-2B fusion proteins with mTur and examined their localization. We found that PKC-2::mTur fluorescence was found in the cytoplasm of intestinal cells but was excluded from the nucleus (Fig. 4b). In addition, fluorescence was detected on the apical, basal and lateral intestinal membranes (Fig. 4b). The laterally-localized PKC-2B colocalized with the lateral membrane marker INX-16::Venus (Fig. 4b). FLP-2 containing DCVs adopt a punctate pattern throughout the intestinal cytosol [29]. FLP-2::mTur containing DCVs are also localized to the basal and lateral intestinal membranes (Fig. 4c), and the laterally localized FLP-2 colocalized with the lateral membrane marker INX-16 (Fig. 4c). Since the basal and lateral intestinal membranes are in contact with the pseudocoelom [46], the localization of both PKC-2 and FLP-2 to the basolateral membranes is consistent with the idea that PKC-2-regulated exocytosis of FLP-2 results in the release of FLP-2 into the pseudocoelom where it could carry out its antioxidant functions by regulating AIY.

## Discussion

This study reveals a previously unreported function for vitamin B5 in promoting organism-wide protection against toxicity caused by oxidative stress (Fig. 1) as well as function for vitamin B5 in selectively and rapidly positively regulating the secretion of the antioxidant neuropeptide-like protein FLP-2 from the intestine (Fig. 2). Vitamin B5 is taken up into cells by the multivitamin sodium dependent transporter SMVT-1 (Fig. 1d), which is required for the effects of vitamin B5 on survival and on neuropeptide secretion (Figs. 1f and 2e). Since the protective effects of vitamin B5 require *flp-2* signaling (Fig. 1e), we propose that vitamin B5 obtained from bacteria is taken up into intestinal cells where it positively regulates FLP-2 secretion from the intestine. Many of the bacterial macronutrients and metabolic byproducts that regulate peptidergic signaling function by activating GPCRs or other receptors on the surface of intestinal cells [22, 51, 52]. By contrast, this study defines a role for vitamin B as an intracellular signal within the intestine, since its uptake is required for its effects.

We speculate that differences in vitamin B5 levels in the bacteria *C. elegans* consumes in its natural environment could regulate intestinal FLP-2 secretion. Indeed, vitamin B5 production can vary widely between different bacterial species and even within species depending on variations the expression of the pantothenate biosynthetic gene cluster [53, 54]. Our previous findings show that once released into the pseudocoelom, FLP-2 functions as an antioxidant signal by stimulating the secretion of a second neuropeptide like protein FLP-1 from the interneuron AIY. FLP-1, in turn, stimulates the SKN-1/Nrf2 antioxidant response in intestinal cells thereby conferring resistance to oxidative stress [28, 29] (Fig. 2a). Thus, dietary vitamin B5-regulated FLP-2 secretion may allow *C. elegans* to rapidly fine-tune the antioxidant response as they consume different bacteria in a constantly changing environment. CoA produced by dietary vitamin B5 is critical for an inter-tissue signaling pathway in which CoA produced in the kidney regulates gut homeostasis in *Drosophila* [55], reinforcing the idea that vitamin B5 uptake can play a critical role in inter-tissue metabolic coordination. Whether vitamin B5 functions specifically in the oxidative stress pathway in *C. elegans* or whether it may function more broadly by regulating the secretion of other signals or by modifying other stress response pathways remains an open question.

We identified two genes that share similarity to mammalian SMVT, *smvt-1* and *slc-5A12*, and our genetic analysis reveals that *smvt-1* and *slc-5A12* appear to function partially redundantly, with *smvt-1* being the primary transporter in the intestine, but *slc-5A12* becoming critical in the absence of *smvt-1* (Fig. 1e). Although *smvt-1* mutants have significantly lower vitamin B5 levels compared to wild type controls, *smvt-1* mutants exhibit superficially normal developmental rates, movement and fecundity (not shown) and they exhibit normal stress tolerance and neuropeptide secretion (Figs. 1e and 2d), suggesting that at least under normal laboratory conditions, *C. elegans* can tolerate significant reductions in vitamin B5 levels without noticeably altered physiology. This is consistent with there being very few reported cases of vitamin B5 deficiency in human populations, although localized deficiencies in certain brain regions are associated with neurodegenerative disease [9, 10].

Mammalian SMVT transports at least three metabolites in addition to vitamin B5: biotin (vitamin B7), lipoic acid, and iodide. Vitamin B5 can competitively inhibit the uptake of these metabolites by SMVT [56]. Thus, it is possible that the effects of exogenous vitamin B5 on stress tolerance and neuropeptide secretion could be consequences of inhibition of uptake of any of these other substrates transported by SMVT-1. However, we do not believe this is the case since *smvt-1* mutants, have normal stress tolerance and neuropeptide secretion in the absence of added vitamin B5 (Figs. 1e and 2d). Instead, we propose that the effects of vitamin B5 exposure on stress tolerance and neuropeptide secretion can be directly attributed to increased cellular vitamin B5 levels.

We found that SMVT-1 localizes to the apical domain as well as the basal and lateral domains in the intestine. This localization pattern differs from that of mammalian SMVT, which is reported to be restricted to the apical domain [43]. We found that SMVT-1 localization and function is dependent upon its PDZ-binding domain (Fig. 3). Mammalian SMVT also requires its PDZ-binding domain for its transport function and for its localization, and the interaction of SMVT with the PDZ domain protein PDZD11 is required for SMVT membrane localization [48]. The *C. elegans* genome does not encode an obvious PDZD11 homolog, but it does encode dozens of PDZ domain-containing proteins that are expressed in the intestine, which may include regulators of membrane localization of SMVT-1.

This study establishes a cell-autonomous function for vitamin B5 a positive regulator of neuropeptide secretion. Since the effects of vitamin B5 exposure on FLP-2 secretion can be detected after as little as 10 minutes (Fig. 2b) vitamin B5 is likely to act via a relatively direct mechanism. B5 itself or one of the four intermediate metabolites that are precursors to CoA biosynthesis are unlikely to regulate exocytosis since the only reported roles for vitamin B5 and its metabolites are as precursors for CoA production. Alternatively, CoA itself or one of the thioester derivatives of CoA such as acyl-CoA, or long chain acyl CoA, could function as regulators of FLP-2 secretion. For example, acyl-CoA is a substrate for ATP production in the mitochondria, and ATP is an essential energy source for vesicle exocytosis and in maintaining ion gradients needed for regulated secretion [57, 58]. Acyl-CoA is also a precursor for phospholipid synthesis, and alteration in phospholipid composition can impact membrane flexibility and affect fusion pore dynamics during exocytosis [59]. Finally, long-chain acyl-CoA is utilized for the palmitoylation of SNAP25, which is required for tethering SNAP25 to membranes for exocytosis [60]. However, the rapid effects of vitamin B5 treatment on neuropeptide secretion that we observe are likely to influence exocytosis at shorter timescales than can be achieved by regulating these metabolic pathways through CoA derivatives.

Instead, we favor a more direct role for CoA itself in promoting FLP-2 secretion. One potential mechanism by which CoA could regulate secretion is through CoAlation. CoAlation involves the covalent attachment of CoA to specific protein cysteine thiols on target proteins, resulting in the alteration of protein activity, subcellular localization and/or conformation in numerous cell types [61]. CoAlation can occur in response to oxidative and metabolic stress and is proposed to protect thiols on proteins from over-oxidation by forming disulfide bonds with cysteine residues that have been oxidized by hydrogen peroxide [5, 61]. We previously found that FLP-2 secretion is positively regulated by hydrogen peroxide generated from mitochondrial superoxide dismutase SOD-3 and we identified a number of putative reactive cysteine residues near the PKC-2 catalytic domain that are required for hydrogen peroxide-induced FLP-2 exocytosis [29]. Therefore, CoAlation of these cysteine residues following their oxidation by hydrogen peroxide could potentially regulate PKC-2 function in this pathway. Alternatively, CoAlation may regulate the activity of some other protein(s) required for exocytosis and/or hydrogen peroxide signaling. CoAlation of cysteine 483 of mitochondrial thioredoxin reductase enhances its enzymatic activity and accelerates cellular antioxidant activity through the antioxidant enzyme thioredoxin [62]. We found that the thioredoxin family member, *trx-3*, negatively regulates hydrogen peroxide levels and FLP-2 secretion [29], raising the possibility that CoAlation may regulate thioredoxin signaling during vitamin B5-induced FLP-2 secretion. Determining whether vitamin B5 regulates FLP-2 secretion via CoAlation and which target proteins are involved will be an interesting area of future investigation.

## Acknowledgements

*C. elegans* strains used in this study were provided by the Caenorhabditis Genetics Center (CGC), which is funded by the NIH National Center for Research Resources (NCRR). We thank Whitaker Cohn at the USC Alfred E. Mann Multi-Omics Mass Spectrometry Core for LC-MS/MS analysis. We thank members of the Sieburth lab for critical reading and discussion of the manuscript.

## Author contributions

AC and DS: conceptualization. JQ and AC: data curation, formal analysis, methodology, writing (original draft). JQ and DS: writing (editing and review)

## Literature cited

1. Fung, T.C., C.A. Olson, and E.Y. Hsiao, Interactions between the microbiota, immune and nervous systems in health and disease. Nat Neurosci, 2017. 20(2): p. 145–155.

2. Pinkosky, S.L., et al., Long-chain fatty acyl-CoA esters regulate metabolism via allosteric control of AMPK beta1 isoforms. Nat Metab, 2020. 2(9): p. 873–881.

3. Martin, W.F., Older Than Genes: The Acetyl CoA Pathway and Origins. Front Microbiol, 2020. 11: p. 817.

4. Li, L.O., E.L. Klett, and R.A. Coleman, Acyl-CoA synthesis, lipid metabolism and lipotoxicity. Biochim Biophys Acta, 2010. 1801(3): p. 246–51.

5. Tsuchiya, Y., et al., Protein CoAlation: a redox-regulated protein modification by coenzyme A in mammalian cells. Biochem J, 2017. 474(14): p. 2489–2508.

6. Amon, P. and I. Sanderson, What is the microbiome? Arch Dis Child Educ Pract Ed, 2017. 102(5): p. 257–260.

7. Sanvictores, T. and R.E. Sina, Vitamin B5 (Pantothenic Acid), in StatPearls. 2026: Treasure Island (FL).

8. Karachaliou, C.E. and E. Livaniou, Biotin Homeostasis and Human Disorders: Recent Findings and Perspectives. Int J Mol Sci, 2024. 25(12).

9. Patassini, S., et al., Cerebral Vitamin B5 (D-Pantothenic Acid) Deficiency as a Potential Cause of Metabolic Perturbation and Neurodegeneration in Huntington’s Disease. Metabolites, 2019. 9(6).

10. Xu, J., et al., Cerebral deficiency of vitamin B5 (d-pantothenic acid; pantothenate) as a potentially-reversible cause of neurodegeneration and dementia in sporadic Alzheimer’s disease. Biochem Biophys Res Commun, 2020. 527(3): p. 676–681.

11. Munshi, M.I., S.J. Yao, and C. Ben Mamoun, Redesigning therapies for pantothenate kinase-associated neurodegeneration. J Biol Chem, 2022. 298(3): p. 101577.

12. Schipper, D.A. and H.M. Schipper, Neurodegeneration with Brain Iron Accumulation. Adv Exp Med Biol, 2025. 1480: p. 291–309.

13. Iuso, A., et al., Mutations in PPCS, Encoding Phosphopantothenoylcysteine Synthetase, Cause Autosomal-Recessive Dilated Cardiomyopathy. Am J Hum Genet, 2018. 102(6): p. 1018–1030.

14. Zhou, B., et al., A novel pantothenate kinase gene (PANK2) is defective in Hallervorden-Spatz syndrome. Nat Genet, 2001. 28(4): p. 345–9.

15. Cavestro, C., et al., Inherited Disorders of Coenzyme A Biosynthesis: Models, Mechanisms, and Treatments. Int J Mol Sci, 2023. 24(6).

16. Rodrigues, D.T., et al., The Caenorhabditis elegans neuroendocrine system and their modulators: An overview. Mol Cell Endocrinol, 2024. 586: p. 112191.

17. Borre, Y.E., et al., Microbiota and neurodevelopmental windows: implications for brain disorders. Trends Mol Med, 2014. 20(9): p. 509–18.

18. Sharon, G., et al., The Central Nervous System and the Gut Microbiome. Cell, 2016. 167(4): p. 915–932.

19. Sherwin, E., T.G. Dinan, and J.F. Cryan, Recent developments in understanding the role of the gut microbiota in brain health and disease. Ann N Y Acad Sci, 2018. 1420(1): p. 5–25.

20. Zhu, S., et al., The progress of gut microbiome research related to brain disorders. J Neuroinflammation, 2020. 17(1): p. 25.

21. Shreiner, A.B., J.Y. Kao, and V.B. Young, The gut microbiome in health and in disease. Curr Opin Gastroenterol, 2015. 31(1): p. 69–75.

22. Covasa, M., et al., Intestinal Sensing by Gut Microbiota: Targeting Gut Peptides. Front Endocrinol (Lausanne), 2019. 10: p. 82.

23. Leeuwendaal, N.K., J.F. Cryan, and H. Schellekens, Gut peptides and the microbiome: focus on ghrelin. Curr Opin Endocrinol Diabetes Obes, 2021. 28(2): p. 243–252.

24. Wang, H., et al., Neuropeptide secreted from a pacemaker activates neurons to control a rhythmic behavior. Curr Biol, 2013. 23(9): p. 746–54.

25. Lee, K. and E. Mylonakis, An Intestine-Derived Neuropeptide Controls Avoidance Behavior in Caenorhabditis elegans. Cell Rep, 2017. 20(10): p. 2501–2512.

26. Liu, C.C., et al., A homeostatic gut-to-brain insulin antagonist restrains neuronally stimulated fat loss. bioRxiv, 2023.

27. Zhang, Y.P., et al., Intestine-specific removal of DAF-2 nearly doubles lifespan in Caenorhabditis elegans with little fitness cost. Nat Commun, 2022. 13(1): p. 6339.

28. Jia, Q. and D. Sieburth, Mitochondrial hydrogen peroxide positively regulates neuropeptide secretion during diet-induced activation of the oxidative stress response. Nat Commun, 2021. 12(1): p. 2304.

29. Jia, Q., et al., Endogenous hydrogen peroxide positively regulates secretion of a gut-derived peptide in neuroendocrine potentiation of the oxidative stress response in C. elegans. bioRxiv, 2024.

30. Arribere, J.A., et al., Efficient marker-free recovery of custom genetic modifications with CRISPR/Cas9 in Caenorhabditis elegans. Genetics, 2014. 198(3): p. 837–46.

31. Mello, C.C., et al., Efficient gene transfer in C.elegans: extrachromosomal maintenance and integration of transforming sequences. Embo J, 1991. 10(12): p. 3959–70.

32. Nonet, M.L., et al., Caenorhabditis elegans rab-3 mutant synapses exhibit impaired function and are partially depleted of vesicles. J Neurosci, 1997. 17(21): p. 8061–73.

33. Egan, C.R., et al., A gut-to-pharynx/tail switch in embryonic expression of the Caenorhabditis elegans ges-1 gene centers on two GATA sequences. Dev Biol, 1995. 170(2): p. 397–419.

34. Li, L. and A.Y. Zinovyeva, Protein Extract Preparation and Co-immunoprecipitation from Caenorhabditis elegans. J Vis Exp, 2020(159).

35. Paulsen, M.T. and M. Ljungman, The natural toxin juglone causes degradation of p53 and induces rapid H2AX phosphorylation and cell death in human fibroblasts. Toxicol Appl Pharmacol, 2005. 209(1): p. 1–9.

36. Ahmad, T. and Y.J. Suzuki, Juglone in Oxidative Stress and Cell Signaling. Antioxidants (Basel), 2019. 8(4).

37. Sidlauskas, K., et al., 5-Hydroxy-1,4-naphthalenedione exerts anticancer effects on glioma cells through interaction with the mitochondrial electron transport chain. Neurosci Lett, 2017. 639: p. 207–214.

38. Rock, C.O., et al., Pantothenate kinase regulation of the intracellular concentration of coenzyme A. J Biol Chem, 2000. 275(2): p. 1377–83.

39. Taylor, S.R., et al., Molecular topography of an entire nervous system. Cell, 2021. 184(16): p. 4329–4347 e23.

40. Yuan, Y. and L. Chen, Transporters in vitamin uptake and cellular metabolism: impacts on health and disease. Life Metab, 2025. 4(3): p. loaf008.

41. Kreuzaler, P., et al., Vitamin B(5) supports MYC oncogenic metabolism and tumor progression in breast cancer. Nat Metab, 2023. 5(11): p. 1870–1886.

42. Mahoney, T.R., et al., Intestinal signaling to GABAergic neurons regulates a rhythmic behavior in Caenorhabditis elegans. Proc Natl Acad Sci U S A, 2008. 105(42): p. 16350–5.

43. Subramanian, V.S., et al., Membrane targeting and intracellular trafficking of the human sodium-dependent multivitamin transporter in polarized epithelial cells. Am J Physiol Cell Physiol, 2009. 296(4): p. C663–71.

44. Vadlapudi, A.D., R.K. Vadlapatla, and A.K. Mitra, Sodium dependent multivitamin transporter (SMVT): a potential target for drug delivery. Curr Drug Targets, 2012. 13(7): p. 994–1003.

45. Gobel, V., et al., Lumen morphogenesis in C. elegans requires the membrane-cytoskeleton linker erm-1. Dev Cell, 2004. 6(6): p. 865–73.

46. McGhee, J.D., The C. elegans intestine. WormBook, 2007: p. 1–36.

47. Peters, M.A., et al., A calcium wave mediated by gap junctions coordinates a rhythmic behavior in C. elegans. Curr Biol, 2007. 17(18): p. 1601–8.

48. Nabokina, S.M., V.S. Subramanian, and H.M. Said, Association of PDZ-containing protein PDZD11 with the human sodium-dependent multivitamin transporter. Am J Physiol Gastrointest Liver Physiol, 2011. 300(4): p. G561–7.

49. Morgan, A., et al., Regulation of exocytosis by protein kinase C. Biochem Soc Trans, 2005. 33(Pt 6): p. 1341–4.

50. Igumenova, T.I., Dynamics and Membrane Interactions of Protein Kinase C. Biochemistry, 2015. 54(32): p. 4953–68.

51. Jyoti and P. Dey, Mechanisms and implications of the gut microbial modulation of intestinal metabolic processes. NPJ Metab Health Dis, 2025. 3(1): p. 24.

52. Hjorne, A.P., I.M. Modvig, and J.J. Holst, The Sensory Mechanisms of Nutrient-Induced GLP-1 Secretion. Metabolites, 2022. 12(5).

53. Sheppard, S.K., et al., Genome-wide association study identifies vitamin B5 biosynthesis as a host specificity factor in Campylobacter. Proc Natl Acad Sci U S A, 2013. 110(29): p. 11923–7.

54. Yoshii, K., et al., Metabolism of Dietary and Microbial Vitamin B Family in the Regulation of Host Immunity. Front Nutr, 2019. 6: p. 48.

55. Miao, T., et al., Renal coenzyme A (CoA) production from VB5 fuels stem cell proliferation and tumor growth. Nat Commun, 2026. 17(1).

56. Chirapu, S.R., et al., High specificity in response of the sodium-dependent multivitamin transporter to derivatives of pantothenic acid. Curr Top Med Chem, 2013. 13(7): p. 837–42.

57. Vos, M., E. Lauwers, and P. Verstreken, Synaptic mitochondria in synaptic transmission and organization of vesicle pools in health and disease. Front Synaptic Neurosci, 2010. 2: p. 139.

58. Dunn, J. and M.H. Grider, Physiology, Adenosine Triphosphate, in StatPearls. 2023: Treasure Island (FL).

59. Ammar, M.R., et al., Lipids in Regulated Exocytosis: What are They Doing? Front Endocrinol (Lausanne), 2013. 4: p. 125.

60. Zhu, M., et al., SNAP25 undergoes phase separation to facilitate the assembly of the synaptic vesicle fusion machinery. Cell Rep, 2026. 45(2): p. 116983.

61. Brett, C. and I. Gout, The two faces of coenzyme A in cellular biology. Free Radic Biol Med, 2025. 233: p. 162–173.

62. Lin, C.C., et al., Coenzyme A protects against ferroptosis via CoAlation of mitochondrial thioredoxin reductase. J Clin Invest, 2025. 135(19).

